# Structural insights into TRAP association with ribosome-Sec61 complex, and translocon inhibition by a CADA derivative

**DOI:** 10.1101/2022.09.28.509949

**Authors:** Eva Pauwels, Neesha R. Shewakramani, Brent De Wijngaert, Anita Camps, Becky Provinciael, Joren Stroobants, Kai-Uwe Kalies, Enno Hartmann, Piet Maes, Kurt Vermeire, Kalyan Das

## Abstract

During co-translational translocation, the signal peptide of a nascent chain binds Sec61 translocon to initiate protein transport through the ER membrane. Our cryo-EM structure of ribosome-Sec61 shows binding of an ordered heterotetrameric TRranslocon-Associated Protein (TRAP) complex, in which TRAP-γ is anchored at two adjacent positions of 28S rRNA and interacts with ribosomal protein L38 and Sec61α/γ. Four transmembrane helices (TMHs) of TRAP-γ cluster with one C-terminal helix of each α, β, and δ subunits. The seven TMH bundle helps position a crescent-shaped trimeric TRAP–α/β/δ core in the ER lumen, facing the Sec61 channel. Further, our *in vitro* assay establishes the CADA derivative CK147 as a translocon inhibitor. A structure of ribosome-Sec61-CK147 reveals CK147 binding the channel and interacting with the plug helix from the lumenal side. The CK147-resistance mutations surround the inhibitor. These structures help in understanding the TRAP functions and provide a new Sec61 site for designing translocon inhibitors.

**Short Summary:** Cryo-EM structures reveal TRAP binding to ribosome-Sec61 complex, and CK147 inhibiting Sec61 by arresting the plug helix inside the channel.

---

About one-third of all eukaryotic proteins are associated with the secretory system that involves the endoplasmic reticulum (ER), Golgi apparatus, trans-Golgi network, and lysosomes (*1*). To reach their respective final destinations, these proteins enter the secretory pathway at the ER, during which proteins are co-translationally translocated across, or inserted into the ER membrane. Protein co-translational translocation is a multistep process that involves the signal recognition particle (SRP)-dependent targeting of the ribosome nascent chain (RNC) complex from the cytosol to the ER membrane and subsequent transport of a nascent polypeptide chain from the ribosome through the Sec61 translocon channel. The heterotrimeric Sec61 translocon consists of α, β, and γ subunits. The Sec61α subunit is composed of ten transmembrane helices (TMHs) that span the ER membrane from the cytosol to ER lumen. These TMHs define the central channel (or pore), and the structural rearrangement of these TMHs regulates the protein translocation steps. The TMH-bundle contains a movable plug domain at the lumenal side of the translocon and a lateral gate that mediate protein transport across and insertion into the ER membrane, respectively (Fig. 1A). In fact, when the RNC complex binds Sec61, the signal peptide (SP) sequence at the N-terminus of the nascent polypeptide interacts with the lateral gate and triggers conformational changes, including the displacement of the plug domain and opening of the channel to the ER lumen (*2*). Once protein translocation is complete, the signal peptidase and oligosaccharyl-transferase (OST) enzymes process the maturation of the translocated preprotein in ER lumen by SP cleavage and N-linked glycosylation, respectively (*3-7*).

**Fig. 1.**
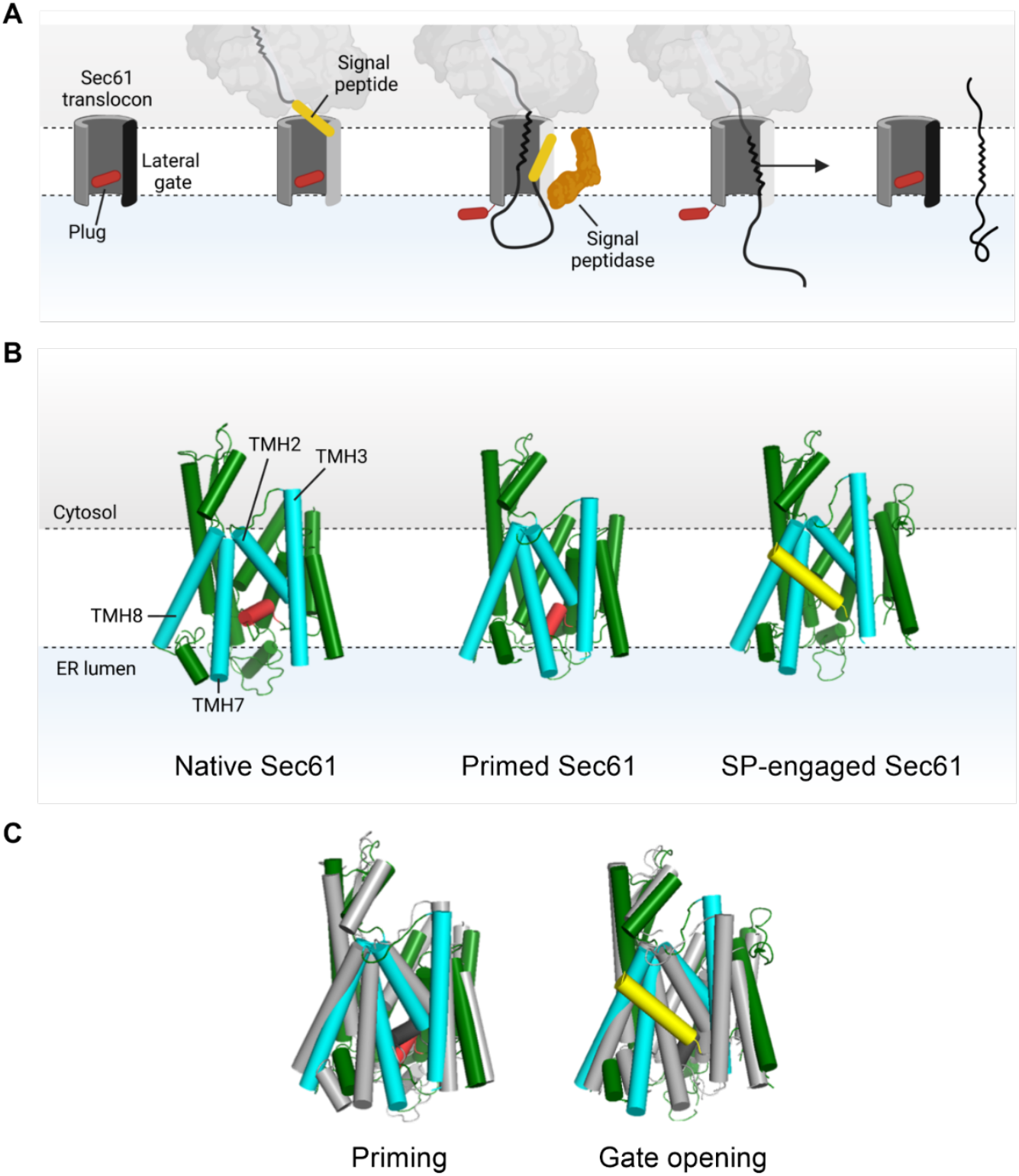
Co-translational translocation across the ER membrane. **(A)** Molecular events in the Sec61 translocon channel during the translocation process. The SP (depicted in yellow) interacts with the lateral gate (depicted in black) of the translocon. This induces the rearrangement of the TMHs of the lateral gate and the plug domain to allow for protein translocation. After SP cleavage by the signal peptidase complex (depicted in brown), the TMD is inserted into the lipid bilayer via opening of the lateral gate, whereas the extracellular part of the protein is translocated to the ER lumen. **(B)** Simplified overview of the different structural states of the Sec61 translocon. Left: Native/closed Sec61 (PDB 5A6U (*8*)). Middle: Ribosome-primed Sec61 (PDB 3J7Q (*9*)). Right: SP-engaged/opened translocon (PDB 3JC2 (*2*)). The Sec61α subunit is shown in green, with the TMHs that form the lateral gate (TMH2, TMH3, TMH7 and TMH8) highlighted in cyan. The plug domain of Sec61α is shown in red. The SP in the engaged/opened translocon (right) is depicted in yellow. **(C)** Comparison of the different structural states. Left: superposition of the ribosome-primed Sec61 translocon (green) with the native translocon (grey). Right: Superposition of the SP-engaged translocon (green) with the ribosome-primed translocon (grey).

Structures representing different functional states of the protein translocation process in the Sec61 translocon (*i*.*e*., closed, ribosome-primed, and SP-engaged/opened configuration; Fig. 1B) have been determined by single-particle cryo-EM in the recent past (*2, 8-11*). Comparisons of these structures revealed the dynamics characteristic of the lateral gate by the repositioning of TMH2 and TMH3 relative to TMH7 and TMH8 during the translocon gating and the repositioning of the plug domain for protein translocation (Fig. 1C). Furthermore, high-resolution cryo-EM structures are available for several accessory proteins such as SRP, the signal peptidase complex, and OST (*3, 5, 6, 11, 12*). Other accessory translocation proteins, such as the heterotetrametric TRanslocoAssociated Protein (TRAP) complex have been identified for which different functions have been proposed (*13, 14*). TRAP, composed of α, β, γ, and δ (ssr1 – ssr4) subunits, is thought to be facilitating the translocation of substrates with weak (low hydrophobicity) or proline and glycine-rich SPs, by ensuring the opened conformation of Sec61 (*15-19*). Furthermore, TRAP might interfere with the final topology of the protein nascent chain (*17*) and subsequently coordinate with OST for the N-linked glycosylation, which suggests a role of TRAP in the biogenesis of glycoproteins (*20, 21*). The upregulation of TRAP expression during ER stress suggests the involvement of TRAP in the ER-associated degradation (ERAD) of improperly folded proteins from the ER lumen to restore ER homeostasis (*22*). In summary, TRAP is involved in various undetermined functions during protein translocation, maturation, and ER stress. The structural information on TRAP subunit assembly and their interactions with the ribosome-Sec61 translocon complex would facilitate a better understanding of the roles that TRAP play in protein translocation. Interestingly, the TRAP complex is present in our eEF2-bound non-translating ribosome-Sec61 translocon complex; the structure, therefore, implies that TRAP is an integral part of the Sec61 translocon network (*19, 23*).

The protein co-translational translocation through ER membrane is essential for the metabolism and survival of cells. The up-or down-regulation of the process is associated with several diseases including cancer (*24-27*). The inhibition of Sec61-dependent protein co-translational translocation is therefore a valuable therapeutic strategy. Most translocation inhibitors have originated from therapeutic screening programs (*28-30*) and Sec61α subunit mutations confer resistance to translocon inhibitors (*31-36*). In general, the resistance mutations emerge adjacent to the lateral gate and/or the plug domain of Sec61, which indicate overlapping mechanisms of action by existing translocation inhibitors and direct impacts of these mutations in disrupting the inhibitor binding.

Examples of characterized small molecule inhibitors of the Sec61 translocon are cotransin (*37*), mycolactone (*35*), and cyclotriazadisulfonamide (CADA) (*38*). CADA was discovered as an anti-HIV agent (*29*) that inhibits the co-translational translocation of human CD4 (huCD4), a type I membrane protein, via specific and selective interactions with the huCD4 SP (*38*). Furthermore, proteomic studies identified five additional CADA-sensitive proteins, namely, SORT, ERLEC1, PTK7, DNAJC3, and 4-1BB, and established CADA as a substrate-selective Sec61 inhibitor (*39-41*). Several CADA derivatives have been evaluated by structure-activity relationship (SAR) studies (*42-45*) in which the cellular huCD4 receptor expression was assessed. An optimized CADA derivative CK147 demonstrated improved huCD4 down-modulating activity from μM to nM range (*44*).

In this work, we identified that CK147 inhibits Sec61-dependent co-translational translocation of huCD4 *in vitro*. Furthermore, CK147-resistance mutations (D60G, R66G, P83H, V102I, and G126K) are located in the lateral gate and the plug domain of Sec61. Our single-particle cryo-EM structure of the ribosome-Sec61 in complex with CK147 reveals that CK147 binds to a site near the plug domain and induces a partially opened conformation of Sec61. In the apo structure of the ribosome-Sec61 translocon complex, with no CK147, the density map permits the tracing of the TRAP-γ subunit and defines its interactions with the ribosome, Sec61, and the remaining subunits of TRAP. This structural information would enhance the current understanding of TRAP’s involvement in the translocation process.

## RESULTS

### Structures of ribosome-Sec61-TRAP complex

We determined the single-particle cryo-EM structures of a primed ribosome-Sec61 complex in which the Sec61-bound ribosome is in a resting phase, *i*.*e*., no peptide being synthesized. The complex was formed by assembling *Oryctolagus cuniculus* (rabbit) 80S ribosome at the *Canis lupus* (dog) ER membrane, as described in the Materials and Methods. The structure was determined at 2.86 Å resolution (Fig. 2A, fig. S1, A to C, and table S1). The ribosome contains eEF2 that interacts with both the 60S and 40S subunits and stabilizes the complex, as observed earlier (*9*). The Sec61 is well defined by the experimental density and the local resolution varies from 3.5 to 6 Å (fig. S1, D to G). As expected, the Sec61 translocon has adopts the primed conformation in which the plug domain seals the bottom part of the channel near the ER lumen. The density map reveals the binding of a well-ordered TRAP-γ subunit alongside Sec61. Only the 60S ribosome that interacts with both Sec61, and TRAP is modeled into the density; we didn’t include the atomic coordinates for 40S or eEF2 in the final structure. The models of the ribosome structures of rabbit (PDB: 6MTE) (*46*) and wild boar (PDB: 3J7Q) (*9*) were used to fit the 60S part to the experimental density. The available structures (*2, 8, 9*) and an AlphaFold (*47*) models of the Sec61α and γ chains were used to fit into the density; the density for Sec61β is weak compared to the α and γ chains, and therefore the β chain was not modeled. Two models, one for TRAP-γ and the other for TRAP (α, β, and δ) trimer were built using AlphaFold and fitted to the cryo-EM density (Fig. 2B and C, and fig. S1, H to J). The secondary structure elements of TRAP-γ model could be fitted to the density. The crescent-shaped density accounting for TRAP-α/β/δ trimer in ER lumen fit the model unambiguously (fig. S1E), however, the map resolution of ∼12 Å at the region is not high enough for secondary structure tracing. Therefore, the atomic coordinates for the TRAP-α/β/δ trimer have zero occupancy and zero B factor in the deposited PDB coordinate file as this part was not included in the final real-space structure refinement cycles.

**Fig. 2.**
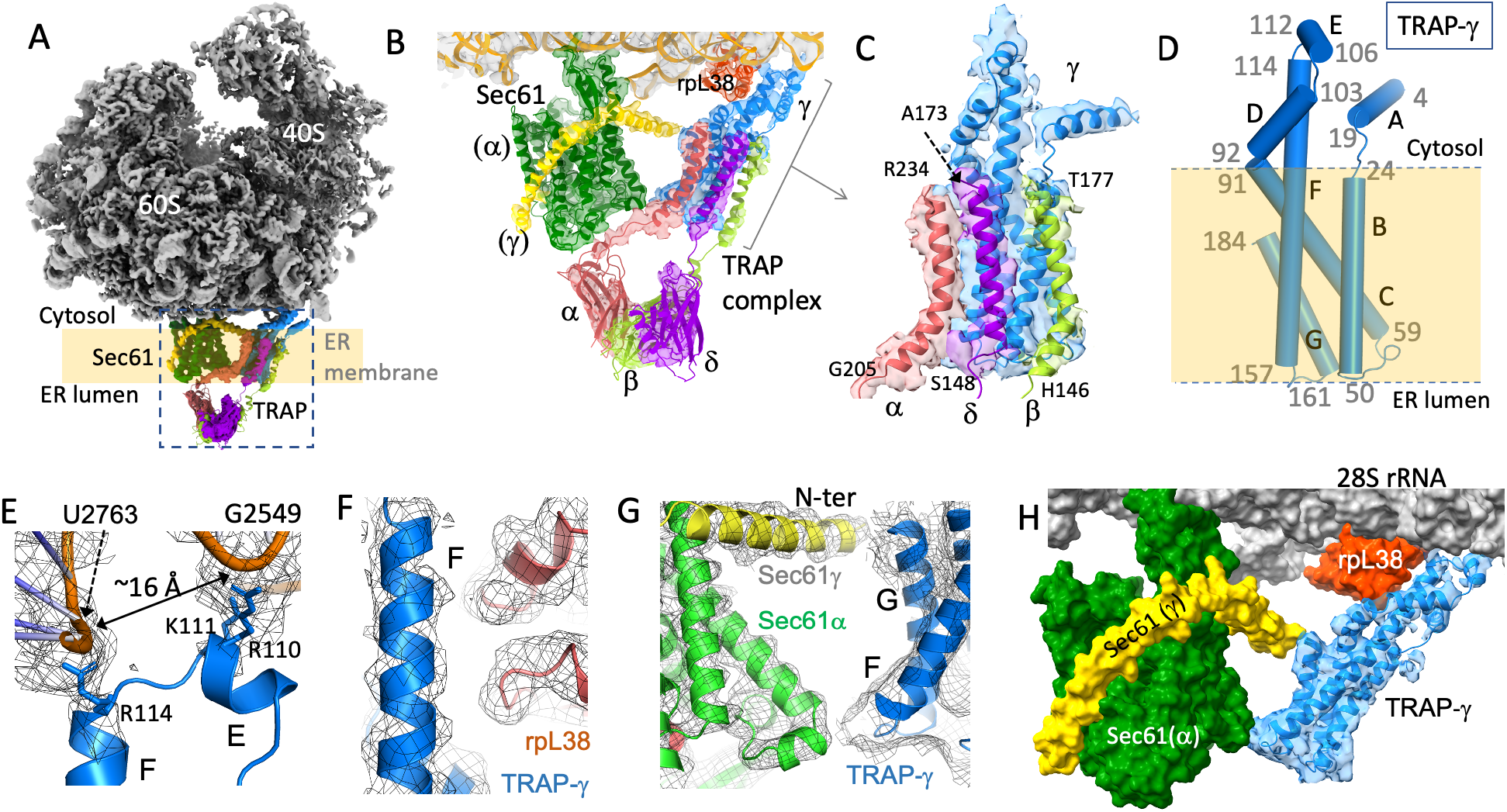
Binding of TRAP to ribosome-Sec61 complex. (**A**) The cryo-EM density map showing the structure of 80S ribosome (gray) -Sec61 (green) -TRAP (heterotetrametric; α, pink; β, light green; γ, blue; δ, purple) complex. The TRAP α. β. and δ subunits are clustered as a trimer in the ER lumen, and the α-chain is positioned near the Sec61 channel. The C-terminal helices, one from each TRAP α. β. and δ–chains cluster with four TMHs of TRAP-γ. (**B**) A zoomed view showing the binding of TRAP and Sec61 to the cryo-EM density map. (**C**) A zoomed view showing the stacking of the TRAP C-terminal helices, one from each α. β. and δ-chains, with TRAP-γ. (**D**) The secondary structure elements of TRAP-γ. **(E)** TRAP-γ interactions with loops of 28S rRNA at two positions that are ∼16 Å apart. **(F)** Relative positioning of the ribosomal protein rpL38 and TRAP-γ. **(G)** Relative positioning of Sec61α and γ subunits with respect to TRAP-γ. (**H)** Molecular assembly of TRAP-γ chain with Sec61 and ribosome.

### Assembly and interactions of TRAP

The structural assembly of the heterotetrametric TRAP and its interactions in the ribosome-translocon complex can help establish the essential yet undefined roles that TRAP plays in ER co- and post-translocation processes (*13-18, 20, 21*). Earlier structural studies both by single-particle (*23*) and cryo-tomography (*13*) have identified the relative positioning of TRAP with respect to the translocon; however, the resolutions of the density maps were not sufficient for tracing the protein chains, which is essential for visualizing the intra- and inter-molecular interactions of TRAP. Our structure (Fig. 2, B and C) enables the fitting of the TRAP-γ chain into the experimental density and identifies its molecular interactions with (i) 28S rRNA and ribosomal protein L38 (rpL38), (ii) α- and γ-chains of Sec61 (Fig. 2B), and (iii) three C-terminal TMHs, one each from the TRAP α, β, and δ chains (Fig. 2C). The TRAP-γ subunit consists of seven helices of which A, D and E are cytosolic, B, C and G are TMHs, and the longest helix F has both cytosolic and transmembrane parts (Fig. 2D). The transmembrane part of TRAP-γ and one C-terminal helix from each TRAP α, β, and δ chains form a seven TMH bundle. The helices B and F of TRAP-γ interface with the TRAP α, β, and δ helices (fig. S1, H to J). The N-terminal parts of the TRAP α, β, and δ chains are clustered as a heterotrimer, which is positioned adjacent to the Sec61 exit channel in the ER lumen (fig. S1K), presumably to interact with the nascent polypeptide chain and/or with other accessory proteins such as OST, as shown earlier (*13*). The TRAP-α subunit is positioned at the Sec61 channel (Fig. 2B and fig.S1K) and thereby TRAP-α is expected to have direct contact with nascent proteins protruding from the translocon, which agrees with the finding that TRAP-α is involved in protein biogenesis (*48*).

TRAP-γ is responsible to place the TRAP complex at its functional site with respect to the translocon (Fig. 2, B and C). The positively charged TRAP-γ residues R110 and K111 of helix E, and R114 of helix F interact specifically with the sugar-phosphate backbone of the 28S rRNA nucleotides U2763 and C2548-G2549, respectively. The RNA backbones at two interacting sites are ∼16 Å apart (Fig. 2E). We hypothesize that the relative positioning of the two positively charged sites of TRAP-γ helps position the TRAP complex in the vicinity of the translocon; i.e., the structure of TRAP-γ at that cytosolic part resembles a “Y-shaped” hook anchoring two RNA “cables” separated by a specific distance of ∼16 Å. The F helix acts as the backbone of TRAP-γ extending from its interacting position with 28S rRNA to the lumenal end of the ER membrane, and the F helix is also supported via its interactions with rpL38 in the cytosol (Fig. 2F) and with Sec61α at the lumenal border (Fig. 2G). The C-terminus of TRAP-γ is positioned near the N-terminal of Sec61γ at the cytosolic side of the ER membrane. The series of observed contacts of TRAP-γ support its precise positioning both in the cytosol and ER membrane regions (Fig. 2H) and place the trimeric TRAP-α/β/δ at the Sec61 channel exit in ER lumen (Fig. 2B, fig. S1K).

### The Sec61α subunit is the target for CK147

In line with a previous report (*44*), CK147 (Fig. 3A) exerts a stronger huCD4 down-modulating effect than the lead CADA compound in transiently transfected HEK293T cells (Fig. 3B), and inhibits the co-translational translocation of huCD4 in a cell free *in vitro* translation assay (Fig. 3C; reduced levels of SP-cleaved huCD4 species; open arrowhead, and fig. S2). Phenotypic analysis revealed significantly higher cytotoxicity of CK147, with a cytotoxic concentration (CC_50_) of 0.36 μM, whereas CADA hardly affects cell proliferation even at 50 μM concentration (Fig. 3D). Furthermore, CK147 affected the cellular expression of CADA clients more profoundly, and reduced the expression of some CADA-resistant proteins in transfected HEK293T cells (fig. S3). However, CK147 maintained some degree of client-specificity, as evidenced by the unaffected expression of surface receptors (such as CD58 and CD86) on T-lymphoid MT-4 cells (fig. S4).

**Fig. 3.**
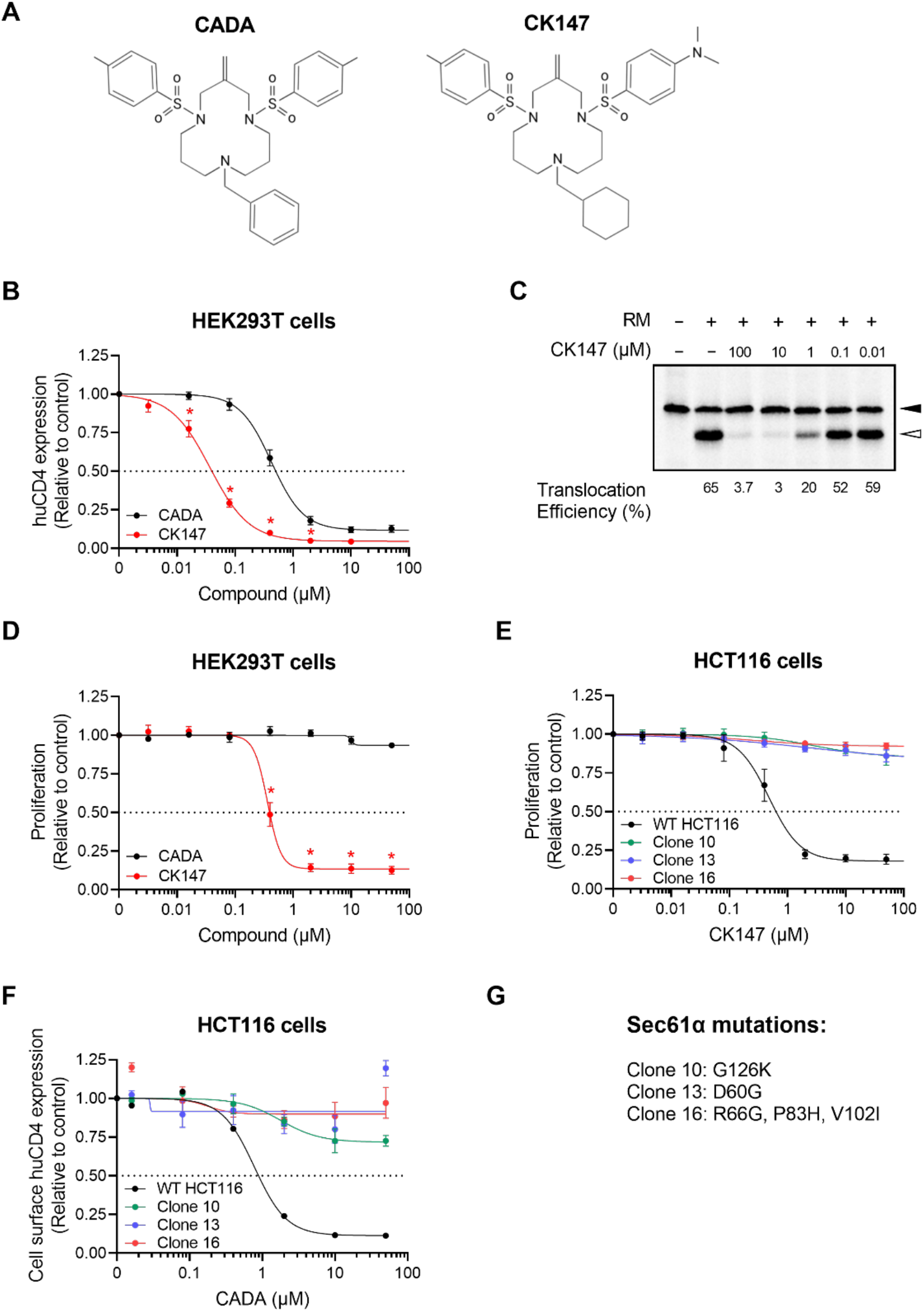
Effect of CK147 on huCD4 expression and cell proliferation in WT and CK147-resistant cells. **(A)** Structures of the lead compound CADA and its unsymmetrical analog CK147 bearing a cyclohexylmethyl tail and a 4-dimethylaminobenzenesulfonyl side arm. **(B)** Concentration-response curves of CADA and CK147 demonstrating the down-modulation of huCD4 in HEK293T cells. Cells were transiently transfected with the huCD4-tGFP-P2A-RFP construct and incubated with different compound concentrations for 24h. Protein levels of tGFP (representing the expression of huCD4) were measured by flow cytometry in compound-treated samples, and were normalized to the DMSO control (set at 1.00). Four-parameter concentration-response curves were fitted to data from three indipendent experiments. Values are mean ± SD; n = 3. Statistical analysis (unpaired t-tests) showed significantly decreased expression of huCD4 when treated with CK147 as compared to CADA (* = p ≤ 0.05). **(C)** Radioblot of the IVT of truncated huCD4 (containing only the D1 and D2 domain). The construct was translated in the presence of RRL, sheep RMs and different concentrations of CK147 or DMSO (1%). Open arrowhead represents the translocated, and thus SP-cleaved, mature huCD4 protein fraction. The solid arrowhead represents the un-cleaved huCD4 preprotein. **(D)** Proliferation of HEK293T cells in the presence of CADA and CK147. Cells were treated for 72h with increasing compound concentrations after which cell proliferation was quantified by measuring the amount of resazurin that was metabolically converted to resorufin within 3h. The data are normalized to the DMSO control (set at 1.00). A four-parameter concentration-response curve was fitted to data from at least three independent experiments. Values are mean ± SD; n = 3. Statistical analysis (unpaired t-tests) showed significantly decreased proliferation of HEK293T cells when treated with CK147 as compared to CADA (* = p ≤ 0.05). **(E)** Same as in (D) but for the proliferation of WT HC116 cells and three CK147-resistant HCT116 clones. Values are mean ± SD; n ≥ 3. **(F)** Cell surface huCD4 expression, measured with flow cytometry, in WT HCT116 and three CK147-resistant HCT116 clones. The different HCT116 cells were transiently transfected with the full-length WT huCD4 construct and incubated with different CADA concentrations for 24h. Cell surface levels of huCD4 in CADA-treated samples were normalized to the DMSO control (set at 1.00). A four-parameter concentration-response curve was fitted to data from at least two replicate experiments. Values are mean ± SD; n ≥ 2. **(G)** Summary of the identified mutations in Sec61α that confer resistance to CK147 in HCT116 cells.

Having established CK147 as a more potent and cytotoxic CADA analog, we next generated CK147-resistant cells to identify potential cellular targets of CADA compounds. Briefly, HCT116 cells were grown in the presence of a toxic concentration of CK147 (25 μM) that exerted maximum anti-proliferative effects in HEK293T cells (Fig. 3D). We selected three CK147- resistat clones (*i*.*e*., clones 10, 13 and 16) for further in-depth phenotypic and genotypic analysis. An untreated control, that went through the same passaging, was also included in our analysis, and is referred to as wild-type (WT) HCT116. Similar to the HEK293T cells, the proliferation of WT HCT116 cells is CK147-sensitive, with a CC_50_ of 0.48 μM (Fig. 3E). In contrast, the selected HCT116 clones showed complete resistance to CK147, as evidenced by the nearly constant proliferation rate at high CK147 concentrations (Fig. 3E; CC_50_ values > 50 μM). In addition, we evaluated the sensitivity of the CK147-resistant cells to the huCD4 down-modulating effect of CADA. As summarized in Fig. 3F, in WT HCT116 cells, the cell surface huCD4 expression is impacted by CADA in a concentration-dependent manner (IC_50_: 0.42 μM), which is in line with the effect seen in HEK293T cells (Fig. 3B) and MT-4 cells (fig. S4), and with previously reported results in other cell lines (*29, 49*). In contrast, the CK147-resistant clones no longer (clones 13 and 16) or hardly (clone 10) responded to CADA’s down-modulatory effect on huCD4 expression (IC_50_ > 50 μM) (Fig. 3F).

Next, the sequence of the Sec61α subunit was analyzed in the resistant clones and aligned to the sequence of WT HCT116 cells. We identified five different heterozygous mutations across the resistant clones, *i*.*e*., D60G, R66G, P83H, V102I and G126K (Fig. 3G). Interestingly, the amino-acid substitutions were all located in one side of the lateral gate, primarily at the interface between TMH2 and TMH3, and in the plug helix. More specifically, in clone 10, the sole amino-acid substitution G126K is located in TMH3. Clone 13 carried a resistance conferring mutation D60G located in the loop connecting the first two TMHs. In contrast to the single mutation in most CK147-resistant clones, the clone 16 has three Sec61α mutations: R66G, P83H, and V102I. The R66G substitution on the plug helix highlights the importance of the positively charged residue for CK147 binding and plug opening activities. The mutation of R66 was also present in the resistant clones of other translocation inhibitors (*31, 32, 35*). The P83H mutation occurs in the TMH2, whereas the V102I mutation is in the connecting loop of lateral gate helices TMH2 and TMH3. The V102I mutation site is at the cytosolic side of Sec61α, and it is the only mutation that is located away from the cluster of the remaining mutations in the channel. To provide a full analysis of the translocon, the Sec61β and γ subunits of the translocon were also sequenced, but did not reveal detectable mutations in any of the three CK147-resistant clones.

### Structure of CK147-bound Sec61 translocon

The experimental density map for the apo ribosome-Sec61 complex, in the absence of an inhibitor, reveals that the Sec61α subunit is in the ribosome-primed conformation with the plug domain sealing the translocon channel (Fig. 4, A and B). The well-defined experimental density, earlier reported structures (PDB IDs 6MTE and 3J7Q) and an AlphaFold generated model helped to reliably model Sec61 and added additional secondary structure features towards the lumenal side of the channel. Here, we numbered the new helices as prime (‘/“) to the preceding TMH number (Fig. 4A). Furthermore, the position of the plug helix in the channel is supported from the lumenal end of the translocon by a newly defined 2-stranded β-sheet (β3–β4; residues 322 – 340), which is located in the middle of the helix H7’. Similarly, we identified two loop structures (residues 196 – 210 and 235 – 241, respectively) that bifurcate the TMH5 and TMH6, respectively, near the lumenal end of the translocon. This β-sheet and the two loops appear to guide the channel opening at the lumenal side of Sec61, as well as the displacement of the plug domain from the translocon channel.

**Fig. 4.**
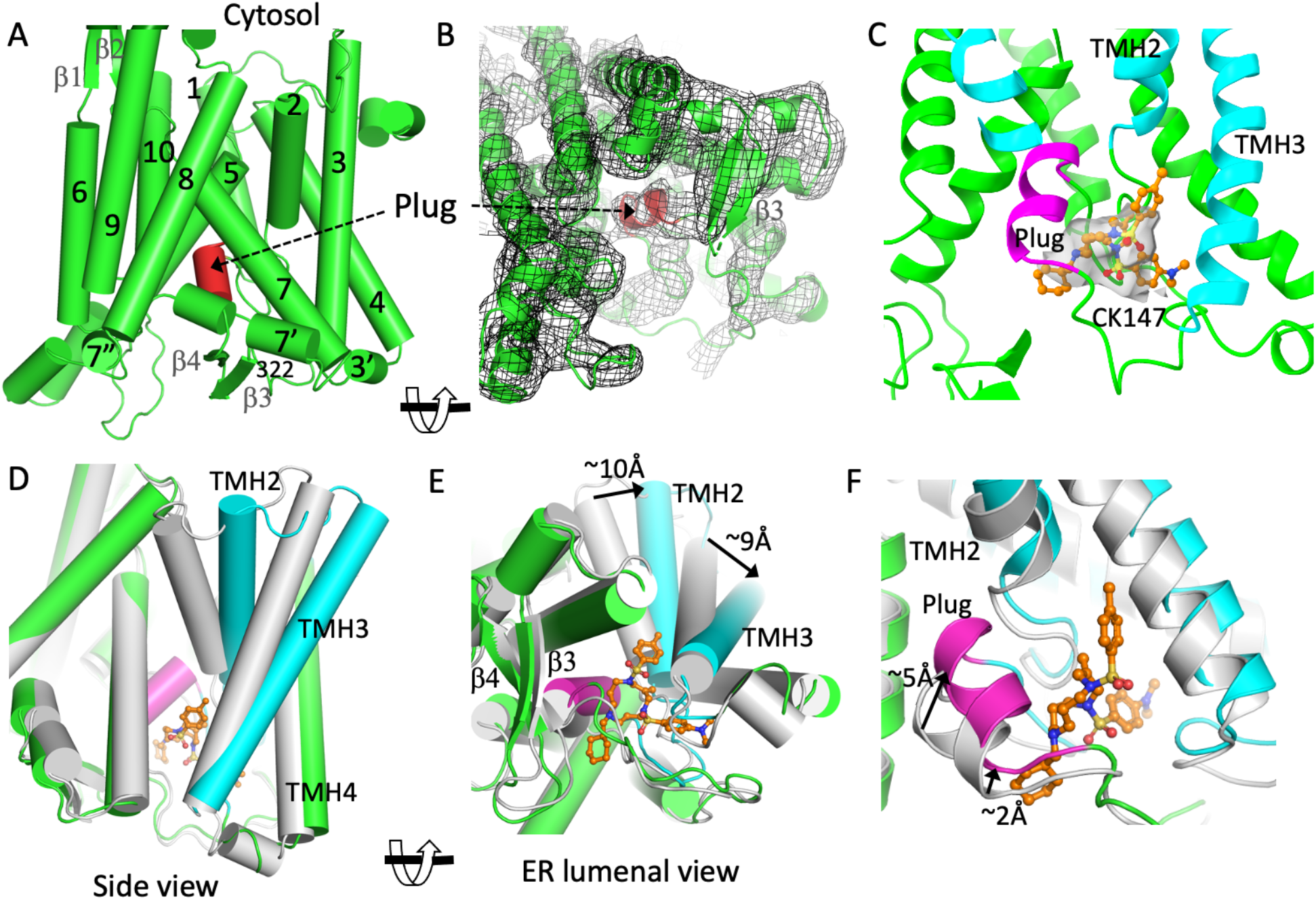
Conformational change of Sec61 upon CK147 binding. (**A)** The Sec61α structure in the ribosome-Sec61-TRAP complex without a bound inhibitor; the plug helix is in red. The transmembrane helices (TMHs) are numbered. The new secondary structural elements modeled into the density are annotated as prime (‘) of the preceding TMHs. (**B)** The ER lumenal view of Sec61 showing the density for different structural elements of Sec61. (**C)** Density for CK147 adjacent to the plug helix (pink) in the ribosome-Sec61-CK147 structure. The docked conformation of CK147 into the site is fitted to the density. (**D)** Superposition of Sec61 structure with CK147 (colored) and without CK147 (gray) reveals the conformational change upon CK147 (orange) binding. (**E)** ER lumenal view of the superimposed structures. (**F)** Repositioning of the plug helix in the pore upon CK147 binding based on the structure superposition in D.

Our second structure is of the ribosome-Sec61-CK147 complex. The inhibitor CK147 was added to the Sec61 translocon before the assembly of ribosome-Sec61 complexes, and the sample was spiked with CK147 (10 μM) prior to the preparation of the grid to ensure high inhibitor occupancy in the complex. Interestingly, a parallel experiment of adding CK147 to a pre-assembled ribosome-Sec61 complex did not result in binding of the inhibitor, and the Sec61α subunit remained in the primed conformation. The cryo-EM data for the CK147-bound complex were also collected on the 200kV Glacios TEM setup and the final density map was calculated at 2.79 Å resolution (table S1). The density for TRAP is significantly weak compared to that in the apo structure (fig. S5A). The experimental density for the Sec61 part confirmed the binding of CK147 deep inside the Sec61α translocon channel (Fig. 4C); the analogous region of the translocon in the apo structure has no such density (Fig. 4B). Superposition of the Sec61 structures with and without CK147 shows significant structural rearrangement of Sec61 upon CK147 binding. Primarily, the lateral gate helices TMH2 and TMH3 are reoriented and TMH4 is shifted as the consequence of CK147 binding (Fig. 4, D and E); the CK147-binding site acts as the pivot point for the observed structural aberrations. In fact, the C-terminal end of the plug helix (Y63 to L69) is shifted by ∼5Å upon CK147 binding (Fig. 4F).

The CK147 binding site is confined to a region between the plug domain and the lumenal end of the Sec61 channel. The local resolution of the density map in that region of the translocon is ∼5 Å (fig. S5, B and C). To ascertain the binding mode and conformation of CK147, primarily for the central 12-member macrolide ring, we applied a molecular docking approach, as discussed in the Materials and Methods section. The final model of the 60S subunit of the ribosome-Sec61-CK147 structure was fitted against the experimental density map by real-space refinement (table S1); the map to model correlation for Sec61α and CK147 are 0.73 and 0.68, respectively.

### CK147 binds near plug helix

The central 12-membered macrolide ring of CK147 is positioned near the plug helix and the compound also interacts with the loops connecting either ends of the plug helix (Fig. 5, A and B). These connecting loops are flexible and in the translocation-competent Sec61 state, they assist the plug domain to move out of the channel towards the lumenal end of the translocon to permit a nascent peptide chain passage through the Sec61 translocon. In the CK147-bound structure, the plug helix and the connecting loops reposition to trap a CK147 molecule at the lumenal side of the translocon. The helices TMH2, TMH3, and TMH4 are rearranged to enclose the inhibitor and participate in CK147 binding. The interactions of CK147 with the residues of Sec61 are shown in a 2D-projection Ligplot in Fig. 5C. In particular, the plug helix backbone and the side chains of residues M65 and R66 interact extensively with the 12-membered core ring of CK147 (Fig. 5D and fig. S5D), whereas the nearby residues A59, D60, P61, L69, S71, and R73 interact from either side (Fig. 5A). Apart from the plug domain, the residue Y131 from TMH3 and M136 from its connecting loop to TMH4 also interact with CK147.

**Fig. 5.**
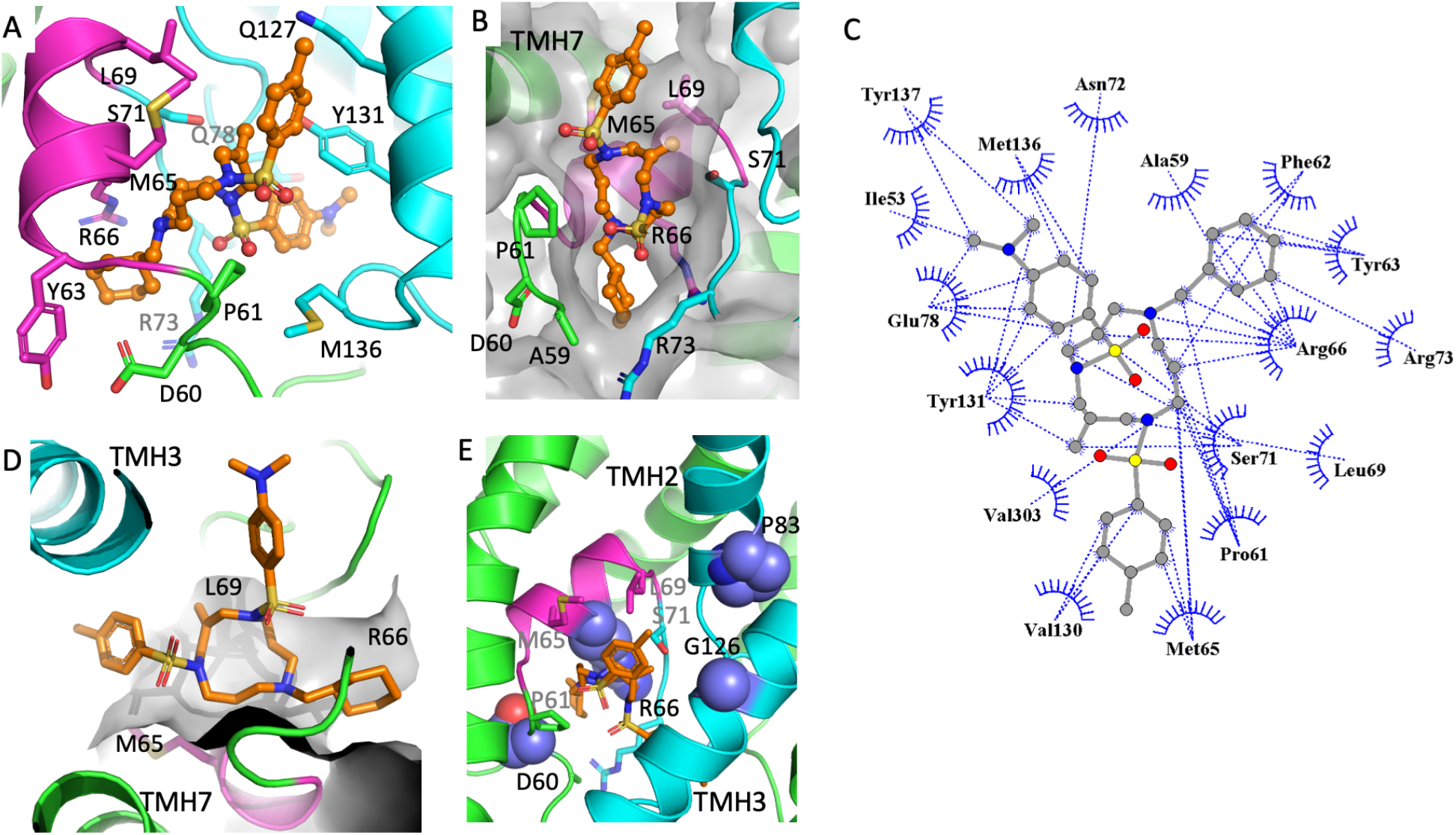
Interactions of CK147 and proximity of resistance mutation sites. (**A)** Mode of binding of CK147 (orange). The interacting Sec61α sidechains are displayed; the plug helix and TMH are in magenta and cyan, respectively. (**B)** Sec61α molecular surface defines the CK147-binding pocket. (**C)** A 2D projection using Ligplot shows the interactions of CK147 with the surrounding residues. (**D**) The proximity and interactions of the plug helix (magenta) with CK147; the interacting surface of plug with CK147 is in gray. (**E**) Four out of five CK147-resistance mutations D60G, R66G, P83H, and G126K surround the inhibitor, whereas the V102I mutation is at the cytosolic side of Sec61.

As described above, the Sec61α mutations D60G, R66G, P83H, V102I, and G126K confer CK147 resistance. Except V102, the remaining mutation sites are in the vicinity of the bound CK147 (Fig. 5E). The residue D60 and the adjacent A59 and P61 are in the loop preceding the plug helix. All three residues interact with the inhibitor, however, P61 interacts more extensively. Apart from the loss of interactions by the D60G mutation, the main-chain flexibility introduced in the region by D60G exchange presumably disfavors the CK147 binding. The residue R66 interacts extensively with CK147 (Fig. 5D), and the significant loss of the inhibitor-protein interactions by R66G mutation is likely to cause the observed CK147 resistance. In fact, only the R66G mutation has been shown to significantly resists mycolactone and restores the translocon activity (*35*), and R66 has the most extensive interactions with CK147 compared to other residues in our structure (fig. S5D). The residue P83 is at the base of TMH2 facing TMH3, and the P83H mutation confer CK147 resistance presumably by rearranging the TMH2 and TMH3 that would not be favorable for CK147 binding. The TMH3 residue G126 is located near CK147, however, with no direct interaction. The introduction of a positively charged long sidechain by G126K mutation might impact the inhibitor entry to the pocket.

## DISCUSSION

### CK147 blocks the plug helix inside the channel

Superposition of the CK147- and SP-bound (PDB ID 3JC2) structures show that the TMHs are rearranged differently in the two complexes (Fig. 6A). A major difference between both structures is that for the SP-bound Sec61 translocon, the plug domain moves out of the channel (Fig. 1), clearing the path for translocation of the protein. In the inhibitor-bound complex, however, the plug domain remains inside the channel, despite the lateral gate TMHs are rearranged. The Sec61 channel in the ribosome-Sec61 complex must switch from the primed to the SP-bound conformation as the signal peptide from the RNC interacts with Sec61. As discussed earlier, our structural study attempt showed that Sec61 does not adapt the inhibitor-bound conformation after the ribosome-Sec61 complex is fully formed, and CK147 should bind Sec61 prior to the docking of ribosomes to Sec61. A previous cell-free *in vitro* translation/translocation study showed that the lead CADA compound is effective at a post-targeting step of huCD4 RNC to Sec61 (*38*). Apparently, a stalled ribosome presenting the huCD4 SP to Sec61 still permits CADA inhibiting the protein translocation. However, no ribosome-Sec61 structure is available for such an early targeting state with the SP approaching a Sec61. Presumably, the conformational changes triggered by an SP permit CADA binding, and the CK147-bound conformation of Sec61 might be a transient state created for SP binding. Thus, the order of inhibitor addition and/or the features of an RNC play roles in the binding of translocon inhibitors.

**Fig. 6.** The potential mechanism of translocon inhibition by CK147. (**A)** Superposition of SP-bound (blue; PDB Id. 3JC2) and CK147-bound (green/cyan) Sec61α reveals that the lateral gate moves differently in two structures; the signal peptide (SP, yellow) binding moves the plug out of the channel, whereas the plug stays in the pore for CK147 binding. (**B, C**) The electrostatic potential surface of SP-bound Sec61α (**B**) and CK147-bound Sec61α (**C**). The binding of CK147 would lock the Sec61 conformation that is unfavorable for SP binding, and protect the channel. **(D)** A schematic representation of the potential mechanism of inhibition by CK147. The approaching SP (yellow) is unable to displace the plug (magenta) out the channel when CK147 (orange) is bound. The elongated nascent chain cannot translocate to the ER lumen, protrudes from the channel at the ribosome-Sec61 interaction site, and is diverted to the cytosol for degradation by the proteasome.

It is generally accepted that an SP recognizes the plug region of the translocon near the lateral gate. The hydrophobic core of the SP is a crucial determinant to push the plug aside, and to permit the nascent peptide passing through the open channel. The SP-bound conformation of Sec61 (Fig. 6B) secures the channel from all sides and permits the nascent polypeptide passage through the ER membrane. The C-terminal end of SP in the looped nascent chain reaches well below in the channel that requires the displacement of the plug. In the CK147-bound Sec61 conformation, the rearranged lateral gate leaves the peptide channel open, whereas the inhibitor locks the plug in the channel and disfavors a productive positioning of the SP helix (Fig. 6C, and D). The CK147-bound conformation of Sec61 appears to permit the peptide entering the channel, however, blocked at the plug helix (Fig. 6C). Thereby, an RNC can potentially destabilize the binding of a weak translocon inhibitor. In this regard the characteristic R66G mutation would reduce the inhibitor binding affinity and the inhibitor can be pushed out to permit SP binding. Also, our lead CADA compound that has different side groups compared to CK147 (Fig. 3A) presumably binds to the site weakly and the binding is reversible (*38, 50, 51*). This interplay between the inhibitor and SP binding could define the client specificity and selectivity for translocon inhibitors. The binding affinity of an inhibitor to Sec61 and the intrinsic binding features of an RNC such as the sequence specificity of an SP for the Sec61 conformational triggering have to be taken into account for finding a protein/SP-specific translocon inhibitor.

### Different inhibitor binding sites in Sec61

The CK147-bound Sec61 structure aligns well with a previously reported structure of the mycolactone-bound Sec61 translocon (*52*). Mycolactone binds at the cytosolic side of the translocon that is ∼20 Å away from the CK147-binding site (Fig. 7A). The local resolution near the mycolactone-binding site in our structure is ∼3 Å (fig. S5C), and no density is observed in that site confirming that CK147 does not bind at or near the proposed mycolactone-binding site. The mycolactone-resistance mutations are primarily observed in the lateral gate and plug domain of the translocon, thus, away from the mycolactone-binding site, but surrounding the CK147-binding pocket (Fig. 7B). Furthermore, mycolactone and CK147 share a 12-membered macrolide core ring, although the ring compositions and side groups differ. A common conformational state of CK147- and mycolactone-bound Sec61, the resemblance in their core 12-membered ring structure, and the overlapping resistance mutation sites in Sec61α suggest that mycolactone might also bind to a second site in which, the core ring would occupy the CK147-binding site, and its long “Southern chain” would extend towards the SP-binding cleft along the mycolactone-resistance mutation sites S82 and T86 in TMH2.

**Fig. 7.**
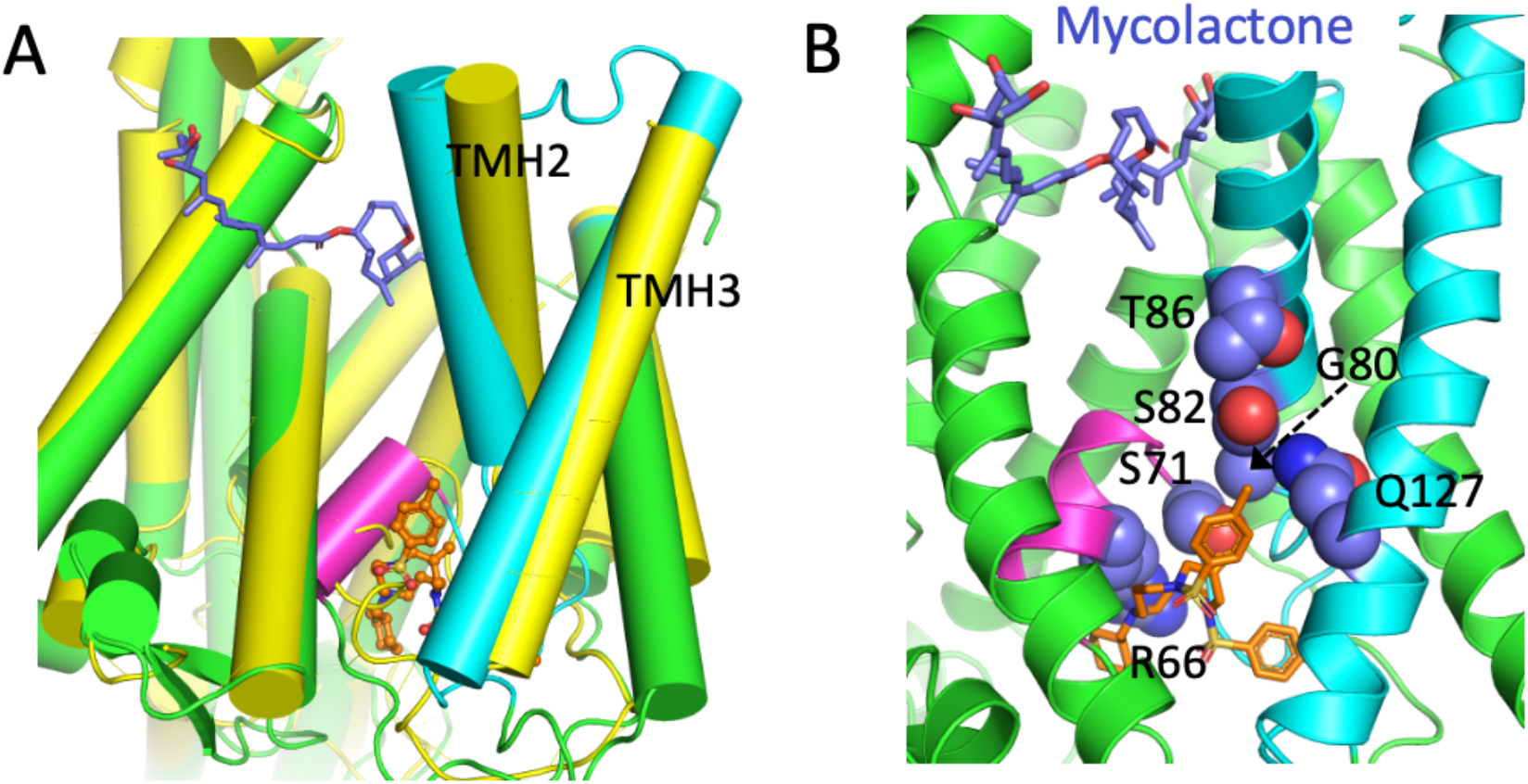
Binding of mycolactone vs. CK147. (**A)** The mycoloctone (blue)-bound Sec61α (yellow; PDB Id. 6Z3T) aligns well with the CK147 (orange)-bound Sec61α (green) suggesting a common mode of action for both inhibitors. (**B)** The key mycolactone-resistance mutation sites (residues with blue spherical atoms) are rather confined to the CK147-binding pocket; the mycolactone is placed based on Sec61α superposition.

A recent cryo-EM study of the Sec61 translocon exposed to a novel cotransin analog, KZR-8445 containing a large cyclic heptadepsipeptide ring, reveals binding of the inhibitor to the SP-binding cleft (*53*). Another recent Sec61 structural study of multiple translocation inhibitors, including mycolactone and CADA, showed an overlapping binding site near the lateral gate of Sec61 for all the inhibitors (*50*). The presented cryo-EM structures, however, were of a post-translocational human/yeast chimeric complex of Sec61/Sec63/Sec71/Sec72 (*50*), and the compounds were soaked at significantly high compound concentrations. In their apo structure, the authors show a preexisting hydrophobic cavity, and Sec61 presumed to be in a non-conventional conformational state, which is different from the native, primed, SP-bound, or CK147/mycolactone-bound state of Sec61 (Fig. 1B and 7A). We did not observe the existence of such a pocket or the presence of an additional density to account for CK147 binding to that region in our structure. Furthermore, the mycolactone and/or CADA-resistance mutation sites in Sec61, such as D60G, R66G and S71F, are distantly located from the proposed preformed hydrophobic cavity. The comparison of structures suggest that inhibitors bind to different sites of Sec61 depending on their shape/size and experimental conditions. Follow up experiments are needed for physiological validation of different binding sites.

### Implications of this study

In this study, we present the binding of a well-resolved TRAP complex that reveals the molecular interactions among the TRAP subunits, and TRAP binding with its partners, *i*.*e*., the ribosome, and the Sec61 translocon. The structure shows that the TRAP complex is a part of the non-translating ribosome-translocon complex, and thereby, reveals TRAP as an integral part of the translocation machinery. The interactions of TRAP with Sec61α and γ subunits complement the earlier findings that TRAP supports the translocation of proteins with less-hydrophobic SPs, and further propose that the protein translocon machinery for the translocation initiation is a multi-protein complex beyond just Sec61. The intra- and inter-molecular interactions of TRAP in cytosol, ER membrane, and ER lumen provide new insights into the organizational and functional roles of TRAP. We expect that this structure provides the framework for experiments such as cross-linking and systematic functional mutation studies to further identify and validate the roles of different TRAP subunits as well as their intra- and inter-molecular interactions at different stages of protein translocation.

Next, we established CK147, an analog of the earlier described CADA compound (*44*), as a potent inhibitor of co-translational translocation. In addition, our resistance study finds Sec61 translocon as the molecular target of CK147. The cryo-EM structure and mutational studies establish the binding of CK147 deep inside the channel, adjacent to the plug helix, and identified a novel Sec61 target site for translocation inhibitor design. The analysis of SAR results on CADA derivatives and other translocon inhibitors in reference to the CK147-binding would help design new translocon inhibitors.

To conclude, this study of a synthetic small molecule, with a 12-membered macrolide ring structure and short side chains, that binds at the plug domain near the lumenal side of the translocon, and potently blocks protein translocation provides several new details for fundamental understanding of co-translational translocation and for inhibitor design. Compared to the recently reported structural studies of Sec61 inhibition, binding of CK147 below the plug domain of Sec61 presents a unique, and unexplored site for translocon inhibitors. The different chemical compositions, and potential binding sites/modes of translocation inhibitors, could pave the way for the structural design and discovery of novel, client-specific inhibitors of the Sec61 translocon for the treatment of human diseases.

## Materials and Methods

### Compounds

Cyclotriazadisulfonamide (CADA) hydrochloride and CK147 were purchased from TCG Lifesciences Pvt. Ltd. (Kolkata, India) who synthesized the compounds following an earlier described protocol (*42, 44*). Compounds were dissolved in dimethyl sulfoxide (DMSO) and stored at a stock concentration of 10 mM at room temperature.

### Plasmids and molecular cloning

The pcDNA3 expression vector encoding WT huCD4, was a kind gift from O. Schwarts (Institut Pasteur, Paris, France). The pEGFP-P2A-RFP expression vector (pEGFP-N1 Clontech backbone) was a kind gift from Prof. Ramanujan Hegde (MRC laboratory of Molecular Biology, Cambridge, UK). The sequence for enhanced GFP (eGFP) was replaced by that of turbo GFP (tGFP). The tGFP-P2A-RFP and huCD4 fragments were cloned into the pcDNA3.1 as described earlier (*54*). Briefly, inserts and expression vectors containing overlapping DNA ends, were amplified using PCR with appropriately designed primers. All PCRs were performed with the Q5 Hot Start HF DNA Polymerase (New England Biolabs, Ipswich, MA, USA) according to manufacturer’s protocol. The PCR products were either purified from the agarose gel, or from the PCR sample using the Nucleospin Gel and PCR clean-up kit (Macherey Nagel, Düren, Germany). Site-directed mutagenesis was performed with the Q5 site-directed mutagenesis kit (New England Biolabs) or NEBuilder HiFi DNA assembly kit (New England Biolabs) following the manufacturer’s instructions. Next, NEB5α competent *E. coli* cells (New England Biolabs) were transformed with the ligated PCR product according to manufacturer’s instructions. Plasmid DNA was isolated using the Nucleospin Plasmid Transfection grade system (Macherey Nagel) supplemented with an endotoxin removal wash. The DNA concentration of all constructs was determined with a NanoDrop 1000 spectrophotometer (Implen, Munich, Germany) and the DNA sequences were confirmed by automated capillary Sanger sequencing (Macrogen Europe).

### Cell-free in vitro translation/translocation assay

The EasyXpress linear template kit (Qiagen, Hilden, Germany) was used to amplify and linearize the DNA of interest from the plasmid using PCR. WT huCD4 contains N-linked glycosylation sites in the extracellular immunoglobulin-like domain D3 and D4 (*55*). To simplify the analysis of the translated products, we generated truncated huCD4 as described previously (*54*). PCR products were purified with the Nucleospin gel and PCR clean-up kit (Macherey Nagel) and transcribed in vitro using T7 RNA polymerase (RiboMAX system; Promega, Madison, WI, USA). RNA was purified from the sample using the Nucleospin RNA clean-up kit (Macherey Nagel). mRNA transcripts were then translated in rabbit reticulocyte lysate (RRL) (Promega) in the presence of L-35S-methionine (PerkinElmer, Waltham, MA, USA). Translations were performed for 20 min at 30°C, in the presence or absence of ovine pancreatic rough microsomes and CK147, supplemented with RNasin (Promega) as described earlier (*56*). Proteinase K (PK) (Roche, Basel, Switzerland) protection assays were performed on ice for 30 minutes and quenched with PMSF (Thermo Fisher Scientific, Waltham, MA, USA). Samples were washed with low-salt buffer (80 mM KOAc, 2 mM Mg(OAc)_2_,50 mM HEPES, pH 7.6) and radiolabeled proteins were isolated by centrifugation (10 minutes at 21,382 g, 4°C). The radio-labelled proteins were then separated with SDS-PAGE on a 4-12% Criterion XT Bis-Tris gels (Bio-Rad, Hercules, CA, USA) in MES buffer (Bio-Rad), detected by phosphor imaging (Cyclone Plus storage phosphor system; Perkin Elmer) and quantified using the ImageQuant software.

### Transient cell transfection and flow cytometry

Cells were seeded in a 6-well plate and incubated overnight to allow adhesion at 37°C prior to transfection the next day. Lipofectamine LTX (Thermo Fisher Scientific) was used for the transient transfection of cells with plasmid DNA, according to the manufacturer’s instructions. Compounds (CADA, CK147or DMSO) were added 6h post transfection. For the cell surface staining of huCD4, cells were collected and washed with ice cold phosphate buffered saline (PBS) (Thermo Fisher Scientific) containing 2% serum (Global Life Sciences Solutions, Traun, Austria). The cells were then stained with a PE-labeled anti-huCD4 antibody (SK3 clone, Cat#344606, BioLegend, San Diego, CA, USA) and incubated in the dark for 45 min at 4°C. Excess antibody was removed by washing the cells with ice cold PBS containing 2% serum. For the flow cytometric analysis of cellular tGFP and RFP levels, no antibody staining was performed. Instead, cells were collected and washed with ice cold PBS. All cells were finally fixed in PBS containing 1% paraformaldehyde (VWR Life Science brand, Radnor, PA, USA) before data acquisition on either a BD FACS Celesta or a BD FACS Fortessa flow cytometer (BD Biosciences, San Jose, CA, USA) running BD FACSDiva 8.0.1 software. All data were analyzed using the software FlowJo X v10 (BD biosciences). To compare the down-modulating activity of CADA and CK147, IC_50_ values were calculated on GraphPad Prism 8 software (San Diego, CA, USA) with a four-parameter concentration–response curve fitting to data from replicate experiments. The absolute IC_50_ value represented the compound concentration that resulted in 50% reduction in the protein level.

### Cell proliferation assay

To analyze the viability of HCT116 or HEK293T cells in presence of compound, a proliferation assay was performed as described by (*33*). Cells are seeded in a black 96-multiwell plate at a density of 25.000 cells/mL After overnight incubation at 37°C, compound is added, and cells are further incubated for 72 hours. Next, the resazurin-based solution Alamar Blue (Thermo Fisher Scientific) is added and the cells are incubated for 3h at 37°C to allow for the metabolic conversion of resazurin to resorufin. Using an excitation wavelength between 530-560 nm, the resorufin emission signal is recorded at 590 nm with Tecan Spark microplate reader (Tecan, Männedorf, Switzerland) and analyzed for calculating the cellular proliferation efficiency upon compound treatment.

### Statistical analysis

Statistical analyses were performed using Graph Pad Prism version 9.3.1. Unpaired t-test with Welch correction was used for the analysis of the flow cytometry data for huCD4 down-modulation in HEK293T cells, and for the proliferation of HEK293T cells in presence of CADA and CK147. Observed differences were considered significant if the calculated p-value was * P ≤ 0.05.

### Selection of CK147-resistant HCT116 clones

HCT116 cells were grown in T25-flasks, treated with 25 μM of CK147 for two weeks and incubated at 37°C. Every two days, the medium was replaced to keep the compound pressure constant. After two weeks, cells that survived compound treatment were collected and diluted over different large petri-dishes to thoroughly dilute the cell suspension. The cells were then further incubated (under CK147 pressure) at 37°C and microscopically monitored. Once cell colonies started to grow, the CK147-resistant cells were isolated by clonal ring expansion, based on a protocol described elsewhere (*33*). Finally, the resistant cells were expanded and cultured in T75-flasks in compound-free medium.

### Library preparation and nanopore sequencing

CK147-resistant HCT116 cells were first incubated with medium containing 25 μM CK147 for 72h at 37°C. Next, RNA was isolated from 5×10^6^ cells using the RNeasy minikit (Qiagen) according to the manufacturer’s protocol, and reverse transcribed into complementary DNA (cDNA) using random primers from the High-Capacity cDNA Reverse transcription kit (Thermo Fisher Scientific). Using specifically designed primers, the Sec61α, β and γ DNA was next PCR amplified with the Q5 polymerase (Promega) according to manufacturer’s protocol. Amplified Sec61α, β and γ fragments were purified using the Nucleospin Gel and PCR clean-up kit (Macherey Nagel). The purified DNA was quantified on a Qubit 4 Fluorometer (ThermoFisher Scientific) and libraries were prepared using the ligation sequencing kit (SQK-LSK 109), from Oxford Nanopore Technologies (ONT), following the manufacturer’s instructions. The amplicons (200 fmol) were first end-repaired with the NEBNext Ultra II End repair kit (New England Biolabs), then purified using AMPure XP beads (Beckman Coulter). Immediately afterward, the end-prepped amplicons were barcoded using the native barcoding expansion kits (EXP-NBD104 and EXP-NBD114, ONT), then the barcoded amplicons were pooled all together and purified using AMPure XP beads (Beckman Coulter). The adapter ligated DNA library was purified with AMPure XP beads, and finally eluted in the elution buffer provided by ONT. DNA library (25 fmol) was loaded on an R9.4.1 flow cell and run for 24 to 48 hours on the GridION sequencing platform. Super accurate base calling, demultiplexing and adapter trimming were done using the GridION built-in MinKNOW software (v21.11). To obtain the consensus sequences, we used Minimap2 (*57*) to align the sequencing reads (single read quality 15 or higher) to the reference, SAMtools v1.9 (*58*) to process the mapping files, and Medaka v1.6.0 (ONT software) to obtain the consensus sequences and variant calling.

### Purification of ribosome-translocon complexes from RRL for cryo-EM

Ribosomes (from RRL), CK147 (10 μM) and dog RMs (0.05 eq/μL) were incubated (in the absence of mRNA) for 45 min at 30°C. Translocon-bound ribosomes were purified from the sample by means of ultracentrifugation as described (*3*). Briefly, membranes were isolated by pelleting the sample through a sucrose cushion (0.5M sucrose, 30 mM HEPES/KOH pH 7.5, 10 mM Mg (OAc)_2_, 500 mM KOAc, 1 mM DTT) for 10 min at 95.029 x g (MLA130 rotor, 4°C). Membrane pellets were then solubilized (30 mM HEPES/KOH, 10 mM Mg(OAc)_2_, 100 mM KOAc, 1.5% digitonin (MP Biomedicals, Santa Ana, CA, USA) and 1mM DTT (Sigma, Saint Louis, MO, USA), pH 7.5) for 1h at 4°C. Next, ribosome-translocon complexes were pelleted through a sucrose cushion (0.5M sucrose, 30 mM HEPES/KOH, 10 mM Mg(OAc)_2_, 100 mM KOAc, 0.3% digitonin, 1 mM DTT, pH 7.5) for 45 min at 560,747 x g (MLA130 rotor, 4°C). The isolated ribosome-translocon complexes were then resuspended (30 mM HEPES/KOH, 10 mM Mg(OAc)_2_, 100 mM KOAc, 0.3% digitonin, 1mM DTT, pH 7.5) and the concentration of the 80S ribosome fraction was determined by measuring the 260 nm absorbance (A260) with a NanoDrop 1000 spectrophotometer. Finally, samples were used in the cryo-EM studies.

### Grid preparation and data collection

Quantifoil® Cu300 R1.2/1.3 + 2nm Carbon (ultra-thin carbon) grids were used to prepare vitreous grids of ribosome-Sec61 complex using a Vitrobot Mark IV. The grids were glow-discharged for 30 s at 15 mA current with the chamber pressure set at 0.30 mBar (PELCO easiGlow; Ted Pella). The glow-discharged grids were mounted in the sample chamber of a Vitrobot Mark IV at 4°C and 95% relative humidity, blotted, and plunge-frozen in liquid ethane at temperature −172°C. Optimized grids were obtained by applying 2.75 μl of the sample (A260 of 19 and 25.5 for the apo and CK147 inhibited sample, respectively), and blotted for 2.5 s using Whatmann grade 2 filter paper with force set to 0. The prepared grids were then clipped and mounted on a 200-kV Glacios TEM (Thermo Fisher Scientific) equipped with autoloader and Falcon 3 direct electron detector as installed in our laboratory. Cryo-EM datasets were collected from the vitrified hydrated grids in counting mode on the Glacios TEM using EPU software version 2.9.0 (Thermo Fisher Scientific). The movies were collected at a nominal magnification of 120,000×, yielding a pixel size of 1.23 Å. Each movie was collected with 40 frames, where each frame received ∼0.8 e/Å^2^, for a total dose of 32 e/Å^2^, and subsequently written as a gain-corrected MRC file. The grids of different ribosome-Sec61 complexes were prepared using the same conditions as described above.

### Data processing

For both structures, the individual movie frames were motion-corrected and aligned using MotionCor2 (*59*) as implemented in the Relion 3.1 package (*60*), and the contrast transfer function (CTF) parameters were estimated by CTFFIND-4 (*61*). The particles were automatically picked using the reference-free Laplacian-of-Gaussian routine in Relion 3.1. The picked particles were initially classified into 2D and 3D classes in cycles to select a set of good particles. The final 3D classification generated a distinct single class of particles. No additional 3D class representing a different structural state of ribosome was detected. The final set of particles for each structure was used to calculate gold-standard auto-refined maps, which were further improved by B polishing and CTF refinement (table S1). Our attempts of multi-body refinement did not improve the quality of the map at the regions of our interest, i.e., Sec61 and TRAP. All data processing were carried out using Relion 3.1.

### Model fitting

The density map for the ribosome-Sec61-TRAP complex was calculated at 2.86 Å resolution (fig. S1). The map revealed the presence of an eEF2-bound ribosome, Sec61 and TRAP. The structure of the rabbit 80S ribosome (PDB: 6MTE) (*62*) was fitted to the 60S part of the density. We did not build the atomic model for eEF2, and the 40S part of ribosome. AlphaFold models generated for Sec61 α and γ chains and the coordinates from a structure of ribosome-Sec61 complex (PDB: 3J7Q) (*9*) were used for building model into the Sec61 part of the density (fig. S1F). Also, two AlphaFold models were generated, one for TRAP-γ and the other for TRAP (α, β. and δ). The local resolution of the map varied from 2.5 Å in the ribosome core to 12 Å in ER lumen (fig. S1E). The TRAP-γ subunit is well-defined in the map (fig. S1G) at a lower counter level than for the core structure, and the AlphaFold model was fitted to the density. The density adjacent to the Sec61 in ER lumen could fit the AlphaFold generated N-terminal trimeric complex of TRAP (α. β. and δ) unambiguously (fig. S1E and K), however, the lower resolution of the density map for the region did not permit the fitting of the heterotrimeric TRAP (α. β. and δ) structure at secondary structure level, and this part was omitted from the real-space structure refinement using Phenix version 1.20.1-4487) (*63*).

The above 60S ribosome structure was fitted to the density map of ribosome-Sec61-CK147 complex. The density for TRAP was present in the map, however, it was poorly ordered. Therefore, no TRAP complex was fitted to this map. The Sec61 conformation altered upon CK147 binding, and the α and γ chains were remodeled into the density in the ribosome-Sec61-CK147 map. The local resolution of the CK147-bound Sec61 varied from 3.5 – 6 Å in resolution, and density accounting for the inhibitor was located adjacent to the plug domain where the map resolution is ∼ 5Å. This region in the apo structure had no density.

### Docking of CK147

While the density in the ribosome-Sec61-CK147 map indicated the position of CK147, the conformation of the inhibitor could not be precisely determined. Therefore, molecular docking was carried out using AutoDock Vina (*64*) as implemented in Chimera v1.15 (*65*) to help find the most probable conformation of CK147. The docking settings were maintained as default and a 25.57 × 28.40 × 18.53 Å^3^ box was set as a cuboid around the region where the density for the inhibitor was found. From top five results based on the docking score, 3 had similar conformations, and the top docked inhibitor model that fits the density with improved conformational state particularly of the 12-membered ring was chosen. The conformation of CK147 from the docking experiment agreed well with the density map. The structure fitted with docked CK147 was refined using Phenix, as described, and the correlation coefficient for CK147 is 0.68 (table S1). The structure figures for the manuscript were produced using ChimeraX v1.2.5 (*66*) and PyMOL v2.4.1 (The PyMOL Molecular Graphics System, Version 2.4.1 Schrödinger, LLC.).

## Acknowledgments

We like to thank Suse Allan for her help with the isolation of sheep and dog microsomes, and Tony Wawina and Bert Vanmechelen for their support to the Nanopore sequencing.

## Author contributions

E.P., K.V. and K.D. designed research; E.P., B.D.W., N.R.S., A.C., B.P., J.S. and K.D. performed research; K.U.K. and E.H. contributed new reagents/analytic tools; E.P., N.R.S, P.M., K.V. and K.D. analyzed data; E.P., K.V. and K.D. wrote the paper.

## Data and materials availability

All data, code, and materials used in the analyses must be available in some form to any researcher for purposes of reproducing or extending the analyses. Include a note explaining any restrictions on materials, such as materials transfer agreements (MTAs). The density maps for the ribosome-Sec61-TRAP and ribosome-Sec61-CK147 structures have been deposited to EMDB under the EMDB Ids. EMD-15860 and EMD-15863, respectively, and the coordinates have been deposited to PDB under the accession numbers 8B5L and 8B6C, respectively. All other data are available in the main text or the supplementary materials.

## Supplementary Materials for

## Supplementary Text

### Cell surface staining and flow cytometry

For the cell surface staining of receptors (Fig. S4), cells were collected and washed with ice cold phosphate buffered saline (PBS, Thermo Fisher Scientific, Waltham, MA, USA) containing 2% serum (Global Life Sciences Solutions, Traun, Austria). Next, cells were stained with antibody for CD4 (SK3 clone, Cat#344606, BioLegend, San Diego, CA, USA), CD25 (Cat#347647, BD Biosciences, San Jose, CA, USA), CD58 (Cat#340295, BD) and CD86 (Cat#555658, BD) and incubated in the dark for 45 min at 4°C. Excess antibody was removed by washing the cells with ice cold PBS (Thermo Fisher Scientific) containing 2% serum (Global Life Sciences Solutions). Cells were then fixed in PBS (Thermo Fisher Scientific) containing 1% paraformaldehyde (VWR Life Science brand) before acquisition on a BD FACS Celesta flow cytometer (BD Biosciences) with BD FACSDiva 8.0.1 software. All data were analyzed in FlowJo X (FlowJo Software), version 10.

**Fig. S1.**
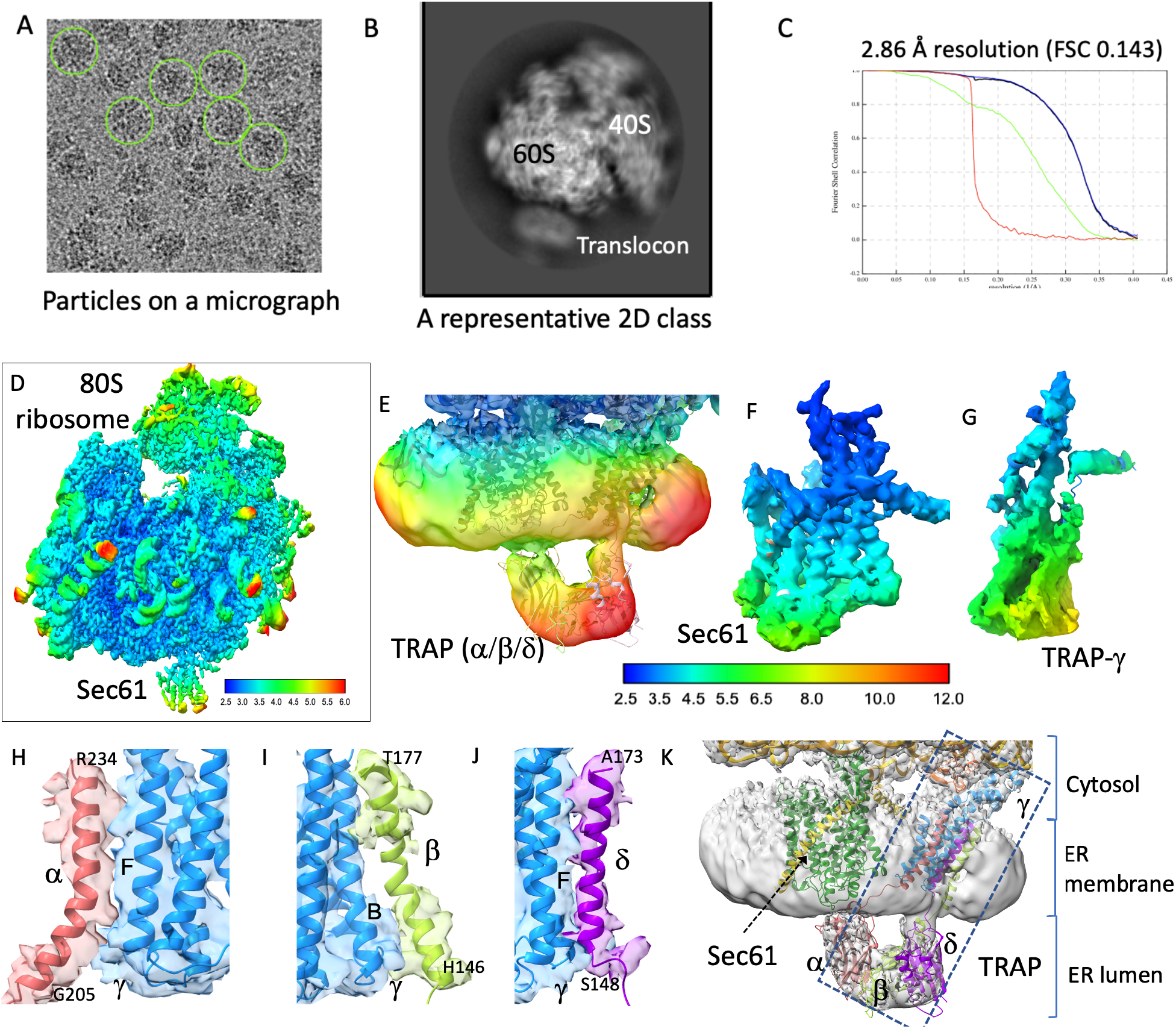
Structure determination and analysis of ribosome-Sec61-TRAP complex. **(A**) A typical micrograph with particles circled. **(B)** A 2D class showing the projection of structural features. **(C)** The overall resolution of the complex is 2.86 Å at FSC 0.143. **(D)** Local resolution of different parts of the structure. **(E)** The local resolution for the TRAP (α, β, and δ) complex located in ER lumen varies between 10 – 12 Å. The local resolution distribution for Sec61 **(F)** and TRAP-**γ (G)**. The experimental density showing the interaction between the transmembrane helices of α and **γ** chains **(H)**, β and **γ** chains **(I)**, and δ and **γ** chains **(J). (K)** The relative positioning of Sec61 and heterotetrametric TRAP in the ER membrane and ER lumen.

**Fig. S2.**
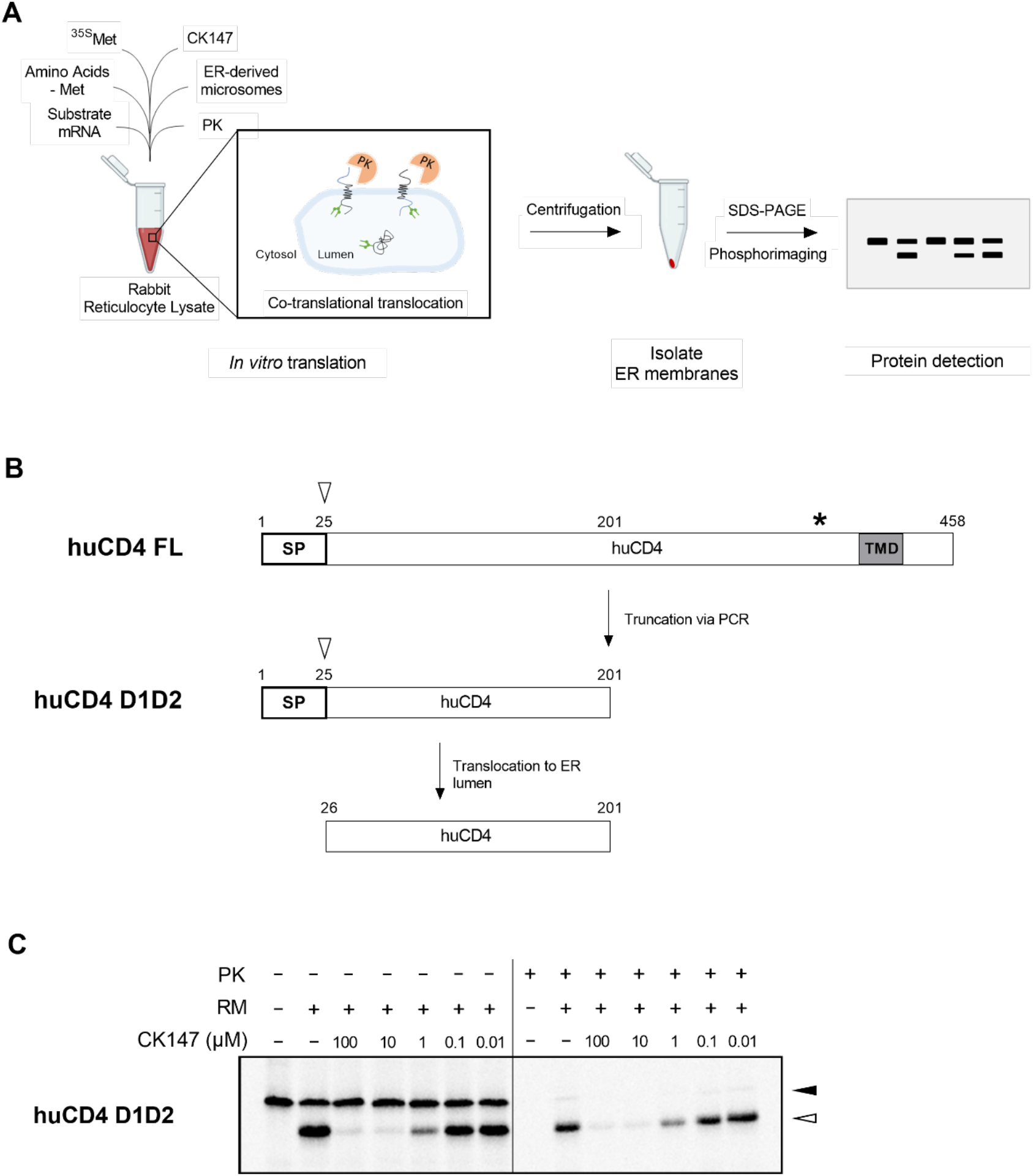
*In vitro* translocation of truncated huCD4 in the presence of CK147 and sheep RMs. **(A)** A schematic representation of the *in vitro* translation/translocation assay. **(B)** A truncated 25 kDa variant of WT huCD4 was used as substrate for the assay, designated huCD4 D1D2, which does not contain the transmembrane region and glycosylation sites. When huCD4 translocates into the ER lumen, the SP is cleaved. **(C)** Same as for Fig. 3C, but with the proteinase K (PK) treatment shown. Protein species that are not translocated, are degraded by the proteinase. As a result, only the translocated, SP-cleaved huCD4 protein is detected on the gel (right side of line). The open arrowhead represents the translocated, and thus SP-cleaved, mature huCD4 protein fraction. The solid arrowhead represents the un-cleaved huCD4 preprotein.

**Fig. S3.**
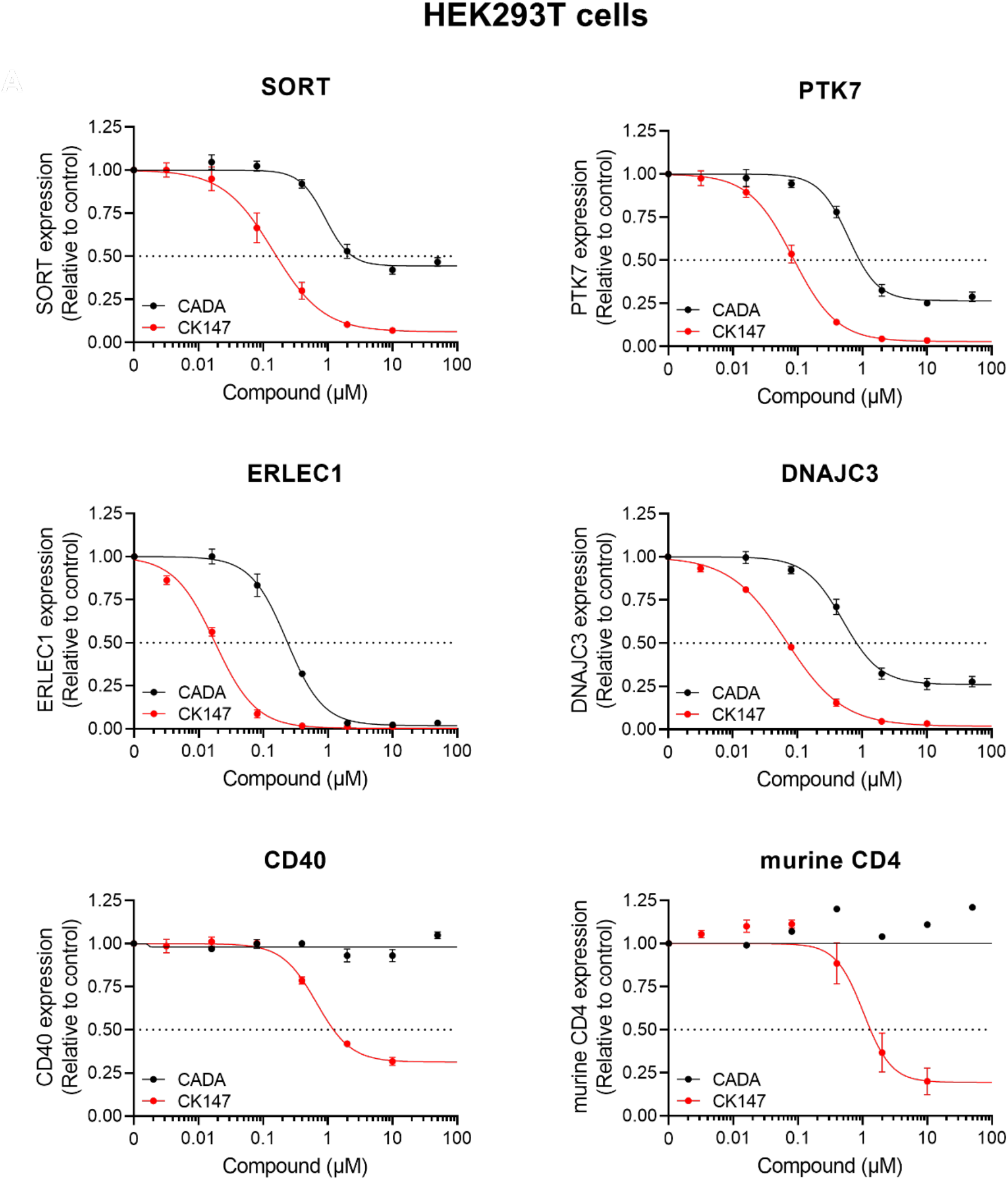
Comparison of CK147 with CADA on protein down-modulation of substrates in transfected HEK293T cells. Same as for Fig 3B. Four-parameter concentration-response curves were fitted to the data from replicate experiments. Values are mean ± SD; n ≥ 3.

**Fig. S4.**
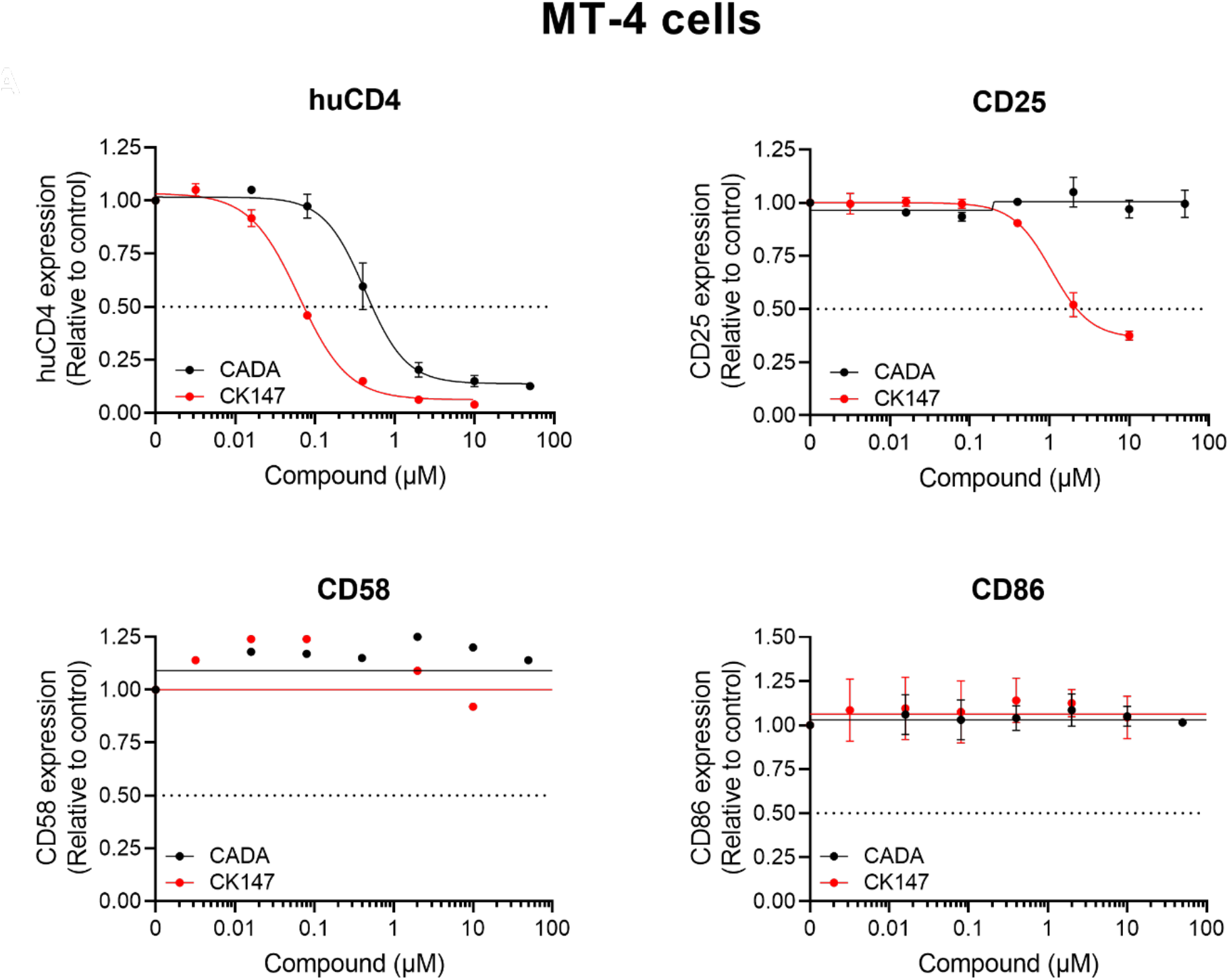
Comparison of CK147 with CADA on protein down-modulation of substrates in CD4^+^ MT-4 cells. Same as for Fig 3B, but in MT-4 cells. Cells were treated with CK147 or CADA for 24h, collected and stained with specific antibodies to measure the cell surface expression of the respective receptors with flow cytometry. On average, 20,000 cells were analyzed for receptor expression to calculate the mean fluorescence intensity. Four-parameter concentration-response curves were fitted to the data.

**Fig. S5.**
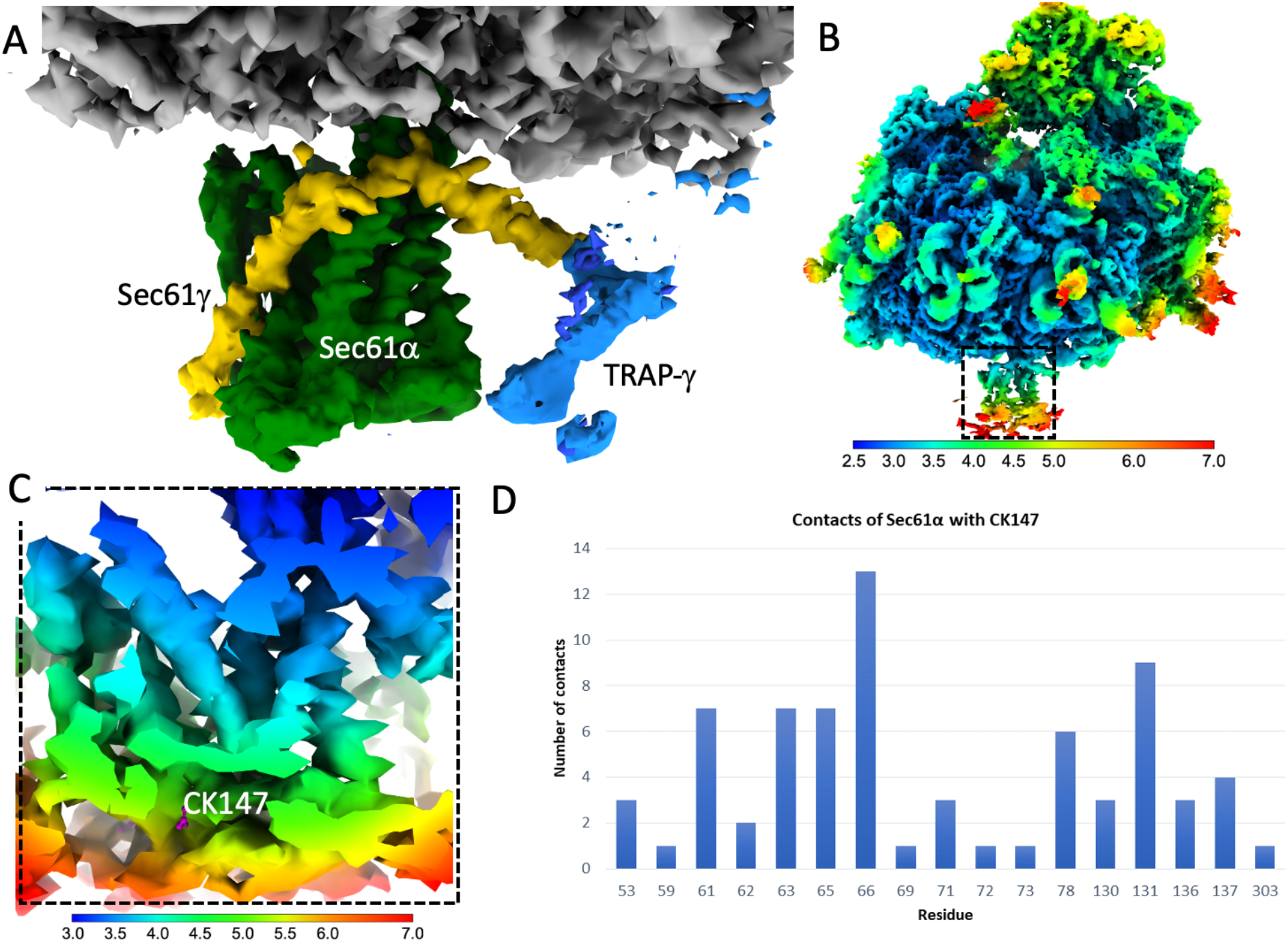
Analysis of the ribosome-Sec61-CK147 structure. (A) The experimental density covering the translocon region of the ribosome (gray) -Sec61 (green) -CK147 structure; the weaker density for TRAP-γ (blue) compared to that in the apo structure suggest that the binding of the translocon inhibitor impacts the positioning of the TRAP complex. **(B)** The local resolution distribution of the ribosome-Sec61-CK147 structure. **(C)** The local resolution distribution for Sec61 varies between 3 - 7Å with the map resolution at the CK147 is ∼5Å. **(D)** The number of contacts of CK147 with Sec61α residues calculated at a maximum cutoff of 4.2 Å. The x-axis indicates the residue number and the y-axis the number of contacts.

**Table S1.** Single-particle cryo-EM data and structure refinement statistics.

|  |  |  |
| --- | --- | --- |
| Structure | Ribosome-Sec61-TRAP | Ribosome-sec61-CK147 |
| PDB ID/EMDB ID | 8B5L/EMD-15860 | 8B6C/EMD-15863 |
| Data collection |  |  |
| Grid type | Quantifoil R1.2/1.3 (2nm C) | Quantifoil R1.2/1.3 (2nm C) |
| Number of grids | 1 | 1 |
| Microscope/detector | Glacios/Falcon 3 | Glacios/Falcon 3 |
| Voltage (kV) | 200 | 200 |
| Magnification | 120,000 x | 120,000 x |
| Recording mode | Counting | Counting |
| Dose (e<sup>-</sup>/Å<sup>2</sup>/frame) | 0.63 | 0.63 |
| Total dose (e<sup>-</sup>/Å<sup>2</sup>) | 32 | 32 |
| Number of frames/movies | 40 | 40 |
| Total exposure time (sec) | 51 | 51 |
| Pixel size (Å) | 1.23 | 1.23 |
| Defocus range (Å) | -6000 to -18000 | -6000 to -18000 |
| Data processing |  |  |
| Number of micrographs used | 2,805 | 1,734 |
| Number of particles picked | 385,028 | 370,000 |
| Particles used for final map | 119,208 | 132,229 |
| Fourier Completeness | 0.923 | 0.924 |
| Map resolution (FSC 0.143; Å) | 2.86 | 2.79 |
| Model fitting |  |  |
| Phenix model resolution (FSC 0.143; Å) | 2.6 | 2.6 |
| Experimental map/model correlation | 0.88 | 0.88 |
| Experimental map/ligand correlation | - | 0.68 |
| Total number of atoms | 143,678 | 138,489 |
| Number of residues/Average B factor (Å<sup>2</sup>) |  |  |
| Protein | 106.72 | 100.03 |
| Nucleic acid | 105.24 | 105.24 |
| Ligand | - | 306.4 |
| Clash score | 9.61 | 9.09 |
| Ramachandran plot; favored/outlier (%) | 96.82/0.03 | 96.63/0.04 |
| Rotamer outlier (%) | 0.47 | 0.50 |
| RMSD bond length (Å)/bond angle (°) | 0.006/1.068 | 0.006/1.079 |
| MolProbity score | 1.70 | 1.70 |

